# Sleep to forget: active control of consolidation and forgetting by slow-wave sleep dynamics

**DOI:** 10.64898/2026.06.15.732460

**Authors:** Ryan Golden, Mingxiao Wei, Samantha Coury, Aviv Mizrahi-Kliger, Karunesh Ganguly, Maxim Bazhenov

**Author notes:** Corresponding author: Maxim Bazhenov.

## Abstract

Sleep supports both the consolidation of new memories and the forgetting of others, but how the cortex flexibly controls these outcomes remains poorly understood. Recent work has shown that two types of Up states may play distinct, competing roles during slow-wave sleep (SWS): slow waves actively consolidate memory traces, whereas delta waves promote their weakening. Here we use a biophysical thalamocortical network model equipped with spike-timing-dependent plasticity to investigate the synaptic mechanisms underlying this dissociation. By manipulating the intrinsic Ca^2+^ dynamics of cortical pyramidal cells, we generate both slow and delta wave Up states within a single network. Using a sequence-learning task paradigm we recapitulate the optogenetic dissociation: removing plasticity during slow waves degrades the memory, while removing it during delta waves enhances consolidation. Mechanistically, the model reveals the longer slow wave Up state affords a spontaneous reactivation phase, occurring after the interfering input, during which the trained memory is selectively reactivated and protected, a phase the truncated delta Up state cannot support. We further find that delta waves sparsen the synaptic representation more than slow waves and predict that the balance between consolidation and forgetting can be flexibly tuned by the ratio of slow to delta waves during SWS.

## INTRODUCTION

The mammalian brain faces a continual dilemma during sleep: it must preserve the neural traces of salient new experiences while simultaneously weakening those deemed less important, all without destabilizing previously stored knowledge (Stickgold, 2005; Born et al., 2006; Poe, 2017). Slow-wave sleep (SWS) has been implicated in both sides of this balance, and a large body of work has established its oscillatory structure as central to memory consolidation (Diekelmann and Born, 2010; Latchoumane et al., 2017; Maingret et al., 2016; Staresina et al., 2015). A widely supported substrate for this process is the reactivation of learning-related neural ensembles during sleep, which reinstates awake activity patterns and is thought to drive their selective strengthening (Wilson and McNaughton, 1994; Ji and Wilson, 2007; Peyrache et al., 2009; Gulati et al., 2014; Ramanathan et al., 2015). Yet how the same machinery could mediate both the protection and the erasure of memory traces has remained unclear.

Part of the answer may lie in a distinction that was historically obscured by the convention of treating all NREM activity with spectral power in the 0.1–4 Hz band as a single SO process. Classic intracellular work distinguished slow (0.1–2 Hz) from delta (2-4 Hz) waves as two separable phenomena with different spatial and temporal properties (Steriade et al., 1993; Steriade and Timofeev, 2003), and subsequent recordings across species have supported the existence of two classes of slow waves during stable SWS (Bernardi et al., 2018; Dang-Vu et al., 2008; Genzel et al., 2014; Mölle et al., 2002; Siclari et al., 2014). Whether these two classes of Up state play distinct functional roles in activity-dependent memory processing, however, was for a long time unresolved.

Recently, Kim et al. (2019) provided a striking causal answer. Using closed-loop optogenetics in rats learning a neuroprosthetic brain-machine interface (BMI) skill, they selectively silenced cortical spiking during the Up states of either slow or delta waves during post-training SWS sleep. Disrupting spiking during slow waves abolished and reversed offline performance gains, whereas disrupting spiking during delta waves enhanced gains beyond control levels. These behavioral effects tracked corresponding changes in spindle nesting and in the strength and persistence of ensemble reactivation, leading the authors to propose that slow and delta waves have dissociable and competing roles: slow waves actively protect and consolidate reactivated memory traces, while delta waves drive their weakening and promote forgetting. The balance between the two could thus bidirectionally set the extent of sleep-dependent consolidation.

This finding reframes the consolidation-versus-forgetting balance as an emergent property of the relative engagement of two Up state regimes, but it leaves the underlying synaptic mechanism unresolved. Because the experiments measured firing rates and ensemble reactivation rather than synaptic weights, they could not specify how reactivation events occurring within a slow wave versus a delta wave Up state come to produce opposite effects on the memory trace, and the authors could only speculate about whether this could be explained by classical activity-dependent mechanisms or if homeostatic downscaling is required (Tononi and Cirelli, 2014). A mechanistic account is needed that explains not only why delta waves might fail to consolidate, but how the cortical network could actively protect a newly formed memory during slow waves while it is simultaneously bombarded with reactivation of competing, task-irrelevant content arriving from the hippocampus.

Here we address this gap using a biophysical thalamocortical network model equipped with spike-timing-dependent plasticity (Gonzalez et al., 2020; Krishnan et al., 2016; Wei et al., 2018). We show that both slow and delta wave Up states can be generated within a single network by manipulating the intrinsic Ca^2+^ dynamics of cortical pyramidal cells, reproducing the shorter Up states characteristic of the ^δ^ regime. Embedding a sequence-learning task in this network and modeling hippocampal sharp-wave-ripple input as interfering “indexing” perturbations time-locked to the Down-to-Up transition (DUt), we reproduce the optogenetic dissociation of Kim et al. (2019): selectively removing plasticity during slow waves degrades the memory, whereas removing it during delta waves enhances consolidation. Critically, the model reveals the mechanism behind this asymmetry—the longer slow wave Up state affords a spontaneous reactivation phase, occurring after the interfering index, during which the trained sequence is selectively reactivated and protected, a phase that the truncated delta wave Up state cannot support. The model further predicts that delta waves sparsen the synaptic representation more than SOs, and that the consolidation–forgetting balance can be flexibly tuned by the ratio of slow to delta waves and the density of novel hippocampal input during sleep.

## RESULTS

### Review of Empirical Phenomenon

Here we briefly review the key results of Kim et al. (2019) in which this phenomenon was initially reported and guided this work. Rats were trained to learn a neuroprosthetic brain-machine interface (BMI) task which monitored neurons in motor cortex, and offline performance gains were assessed across a post-training block of SWS. (Fig 1A). During SWS, Up states were classified as part of either slow or delta waves based on waveform features (Fig 1B). Briefly, an Up state preceded by a large down state deflection peak which crossed the upper threshold is labeled a slow wave, whereas an Up state lacking that preceding large peak is labeled a delta wave. Using this criteria, they detected slow (SO) and delta (δ) waves online during SWS (Fig 1C) and deployed closed-loop optogenetic silencing of cortical spiking, timed to the Up states of one type or another to isolate the consolidation effects. Relative to control optogenetic silencing during slow waves (OPTO_SO_) abolishes and reverses the gain, producing a performance drop, whereas disrupting spiking during delta waves (OPTOδ) enhances the gain beyond control levels.

**Fig 1.**
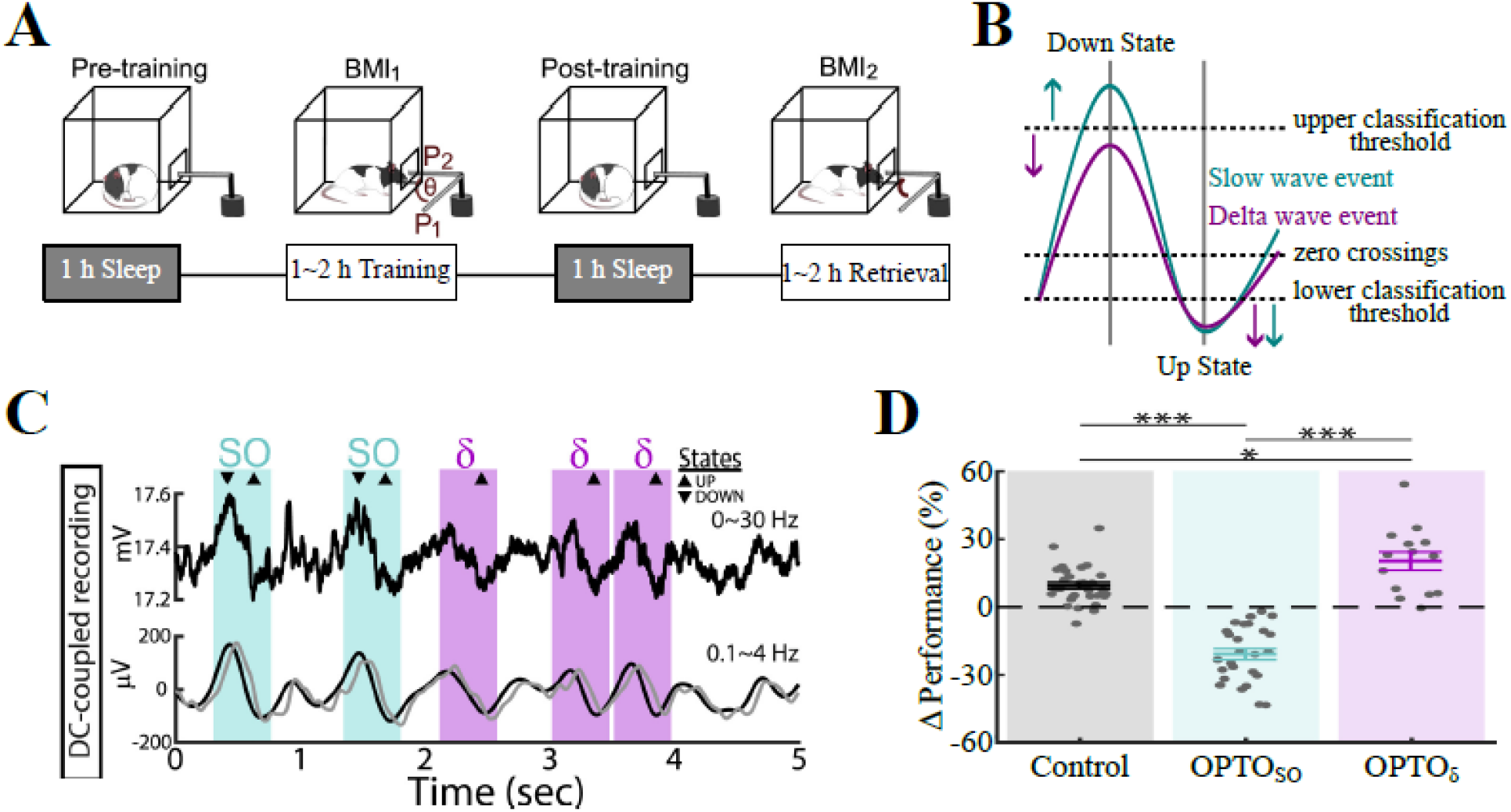
Review of empirical phenomenon. **A**. Schematic of rodent BMI task and sleep consolidation paradigm. **B**. Schematic illustration of operational definition and classification criterion for empirical slow and delta wave events. **C**. Examples of raw and filtered traces with DC-coupled recordings. Traces were filtered using online (gray) and offline (black) digital filters. Slow and delta waves are marked by blue and magenta hereafter. **D**. Task performance changes from BMI_1_ to BMI_2_ in 3 conditions: control (gray), optogenetic silencing of slow waves (OPTO_SO_; blue) and delta waves (OPTOδ; magenta). [Panels A,C, and D modified from Kim et al. 2019]

### Network Model

Throughout the study we utilize a modified version a biophysical thalamocortical model (Wei et al. 2016; Wei et al., 2018; Gonzalez et al., 2020). Briefly (see **Methods** for more detail), the basic thalamocortical circuit (Fig 2A) had a single cortical layer composed of excitatory pyramidal cells (PYs) and fast-spiking, inhibitory interneurons (INs), and a single thalamic layer composed of excitatory thalamocortical cells (TCs) and inhibitory reticular interneurons (REs). All cell types were modeled according to the Hodgkin-Huxley formalism. Nearly all synaptic connections were set deterministically within a local radius which depends on the pre- and post-synaptic cell types, and held constant, except for PY-PY synapses which were subject to plasticity on AMPA receptors. These PY-PY synapses were initialized probabilistically within their local radius, with weight values drawn from a Gaussian distribution (Fig 2B), and were allowed to evolve according to local anti-symmetric spike-timing-dependent plasticity (STDP) rules (Fig 2C). Transitions between awake and slow-wave sleep (SWS) states were simulated by changing the conductance of intrinsic cellular and extrinsic synaptic ion channels to mimic the effects of the neuromodulator tone of each brain state [Krishnan et al., 2016].

**Fig 2.**
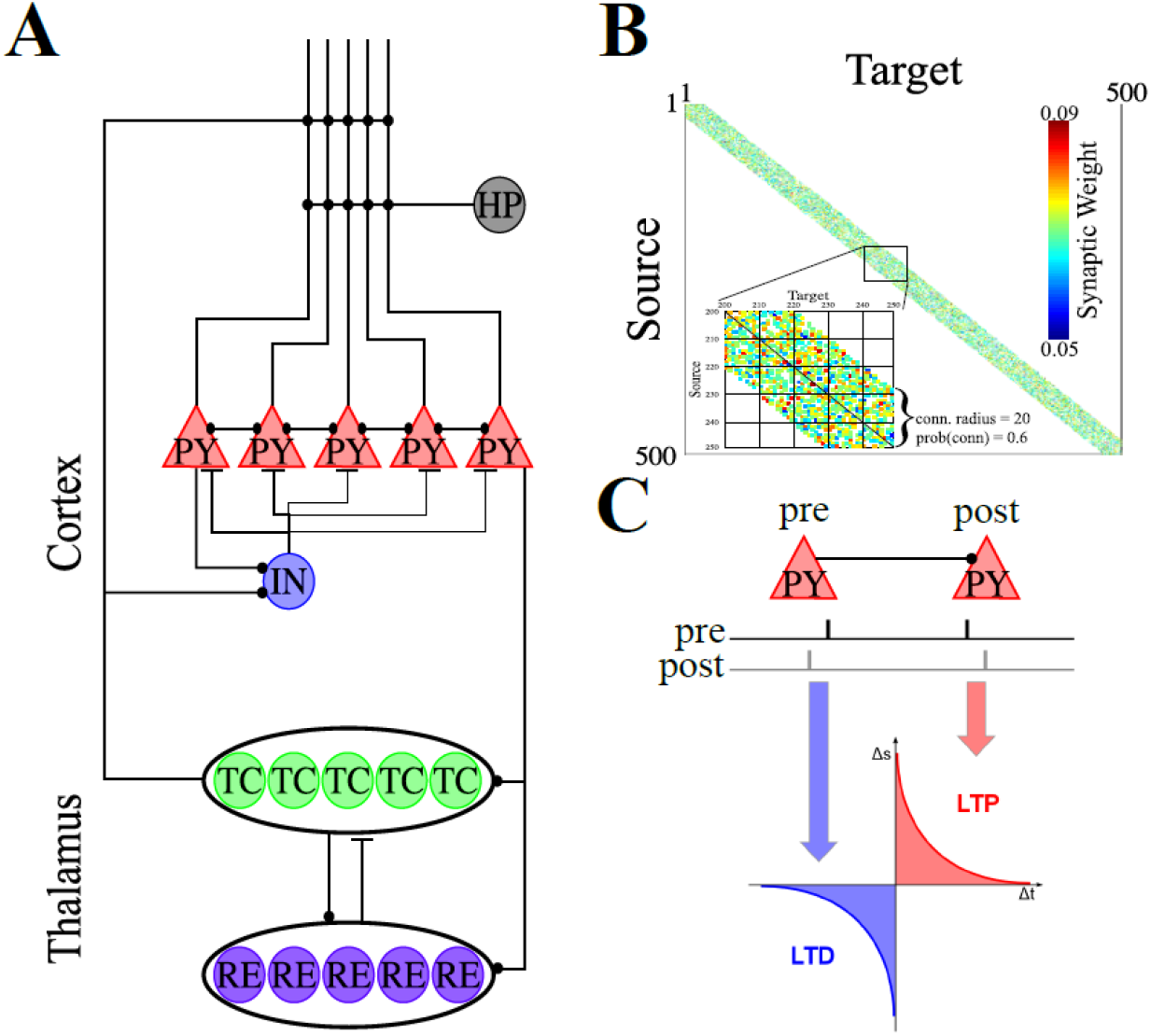
Thalamocortical model architecture. **A**. Basic network architecture (PY: excitatory pyramidal neurons; IN: inhibitory interneurons; TC: excitatory thalamocortical neurons; RE: inhibitory thalamic reticular neurons). Excitatory (inhibitory) synapses are represented by lines terminating in a dot (bar). **B**. Initial weighted synaptic matrix for the PYs. The color represents the strength of the AMPA connections between PY neurons, with white indicating the lack of synaptic connection. The inset shows a zoom-in of the subregion where training occurs (PYs 200-249). **C**. Schematic illustration of the anti-symmetric STDP rule used.

#### Slow and Delta Wave Dynamics

To account for the heterogeneity of Up states seen across both slow and delta wave events, we manipulated the intrinsic calcium dynamics of PYs. Briefly delta waves were produced by increasing the gain of dendritic Ca^2+^ accumulation in PYs, thereby strengthening Ca^2+^-dependent K+ adaptation and shortening Up states (Sanchez-Vives and McCormick, 2000; Bazhenov et al., 2002; Hill and Tononi, 2005; Andrade et al., 2012), while simultaneously accelerating the timescale of spontaneous miniature EPSP release, shortening Down states (Timofeev et al., 2000; Bazhenov et al., 2002). The model was stochastically switched between the slow or delta wave regime every 3 seconds during SWS, with each switch having a 75% chance of transition to a slow wave regime and a 25% chance of transitioning to a delta wave regime. This asymmetry in time spent in regimes was introduced to counter the different event frequencies in the model and keep the total number of Up states roughly equivalent during each regime.

An example of PY activity during both slow and delta wave periods is shown in Fig 3A, where the network begins in a delta wave regime with 5 Up states before transitioning to a slow wave regime for 2 Up states, back into a delta wave regime for 7 Up states, and a single Up state from the slow wave regime at the end. Up and Down states are clearly visible in the simulated LFP (Fig 3B), with spiking from individual PYs locked to the Up states of both regimes (Fig 3C). In the model, Up states from each regime can be clearly identified by filtering the simulated LFP in either the slow wave (0.5-2 Hz; Fig 3D) or delta wave (2-4 Hz; Fig 3E) frequency bands, and all Up states are discernible when the simulated LFP was filtered in the combined frequency band (0.5-4 Hz; Fig 3F).

**Fig 3.**
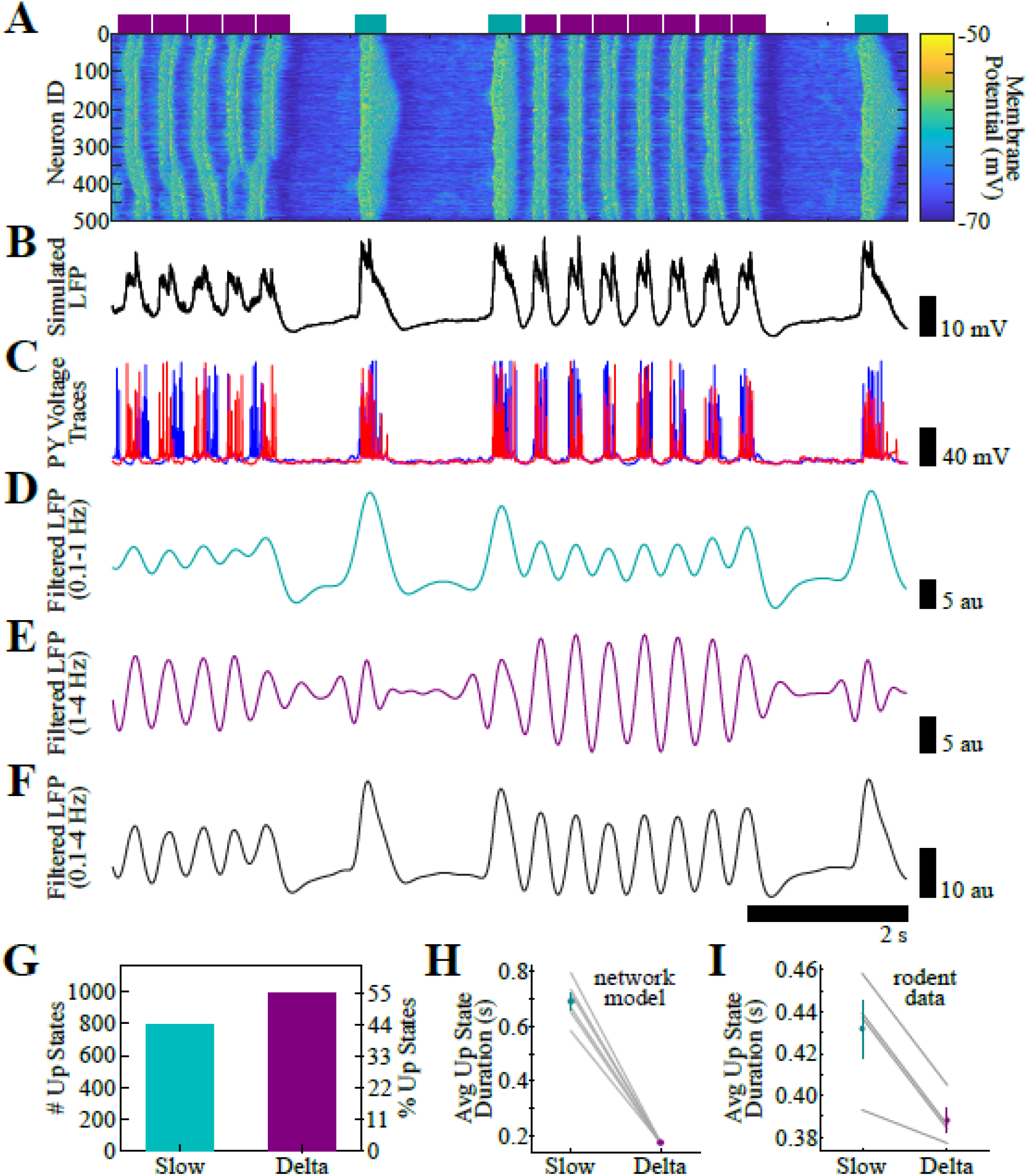
Network dynamics during SWS with a mixture of slow and delta waves. **A**. Membrane potential (color scale) of PYs (y-axis) over time (x-axis) during SWS. **B**. Simulated raw LFP. **C**. Examples of two PYs membrane potential traces. **D**. Filtered LFP in the slow wave (0.1 - 1 Hz) band. **E**. Filtered LFP in the delta wave (1 - 4 Hz) band. **F**. Filtered LFP in the combined slow and delta wave (0.1 - 4 Hz) band. **G**. Bar plot showing the total number of Up states during slow and delta wave regimes during an example simulation. **H**. Mean and SEM of Up state duration across random seeds in the slow (blue) or delta (magenta) regimes in the thalamocortical model. Gray lines show relationship within a given random seed. **I**. Mean and SEM of Up state duration across animals classified as slow (blue) or delta (magenta) regimes in the thalamocortical model. Gray lines show pairwise relationship within a given animal.

The total number of delta (996; 55.4%) and slow wave (801; 44.6%) Up states are shown in Fig 3G, with a slight bias towards delta waves. Finally, in addition to the difference in Up state frequency, the model exhibits substantially shorter duration Up states in the delta wave regime compared to the slow wave regime (Fig 3G), in agreement with new empirical analysis (Fig 1D). Finally, Fig 3H shows the total number of delta (996; 55.4%) and slow wave (801; 44.6%) Up states during SWS in this example simulation.

### Consolidation of Sequence Memory

Next, we simulated a simple sequence learning paradigm to understand how this mixture of slow and delta waves affected memory consolidation (Fig 4A,B). During the awake state, DC current input was provided sequentially to 5 groups of 10 PYs (Fig 4C, left) to force spiking and facilitate encoding of the sequence ABCDE through STDP rules. Recall performance was tested by providing DC current input to the PYs in group A and testing for the extent of pattern completion (Fig 4C, right; see **Methods** for more detail). Finally, during SWS, we also provided DC stimulation to simulate the effects of hippocampal sharp-wave ripple input to cortex carrying other task-irrelevant memory content (Fig 4D). This input was time-locked to the Down-to-Up transition (DUt) of each Up state and stochastically cycled through 4 sequences (ABDEC, BCADE, CAEBD, ACDBE) with equal probability. The boxplots in Fig 4E show that recall performance improved with awake training (T2; Post-Training) relative to at Baseline (T1), and that performance was maintained with this particular slow/delta wave mixture of SWS (T3; Post-Sleep).

**Fig 4.**
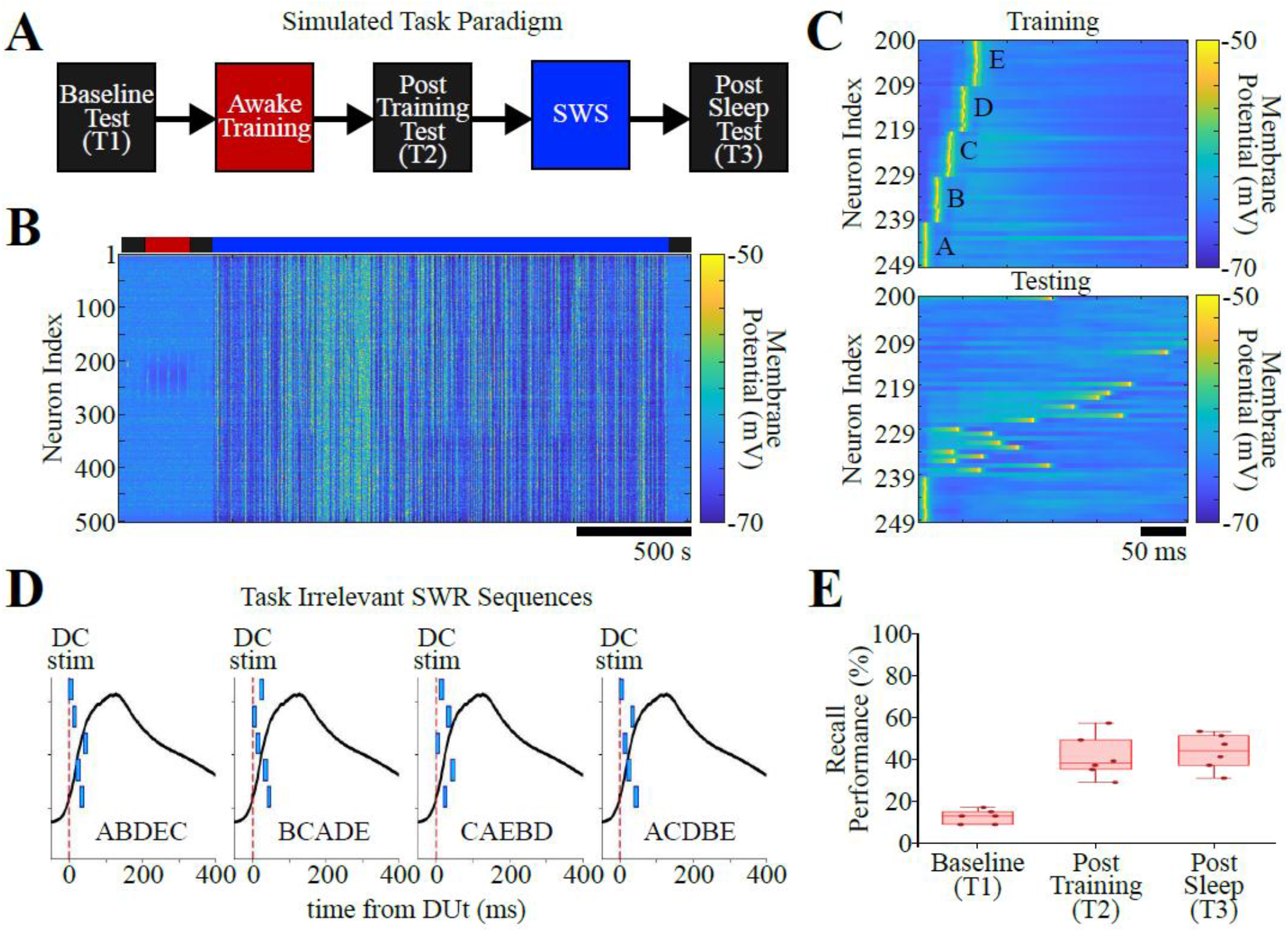
Consolidation of memory sequence during SWS. **A**. Simulated task paradigm awake training of a sequence memory followed by a period of SWS, and 3 testing periods. **B**. Network activity depicting membrane potential (color scale) of 500 PYs on the y-axis over time (x-axis). **C**. Examples of network activity during one bout of sequence training (top), during a single test trial (bottom), where only the first letter was activated, and performance was evaluated by measuring pattern completion. **D**. Schematic illustration of how the 4 task-irrelevant SWR sequences are applied by closed-loop stimulation at the DUt during each Up state of SWS. **E**. Boxplots showing recall performance during each of the 3 test periods across all random seeds.

### Blocking Plasticity during Slow or Delta Waves Recapitulates Optogenetics Findings

Using this consolidation paradigm, we simulated a version the optogenetics experiments (see Fig 1D) reported in the empirical work Kim et al., 2019. We did this by blocking plasticity by turning off STDP during either slow wave (Fig 5A, left) or delta wave (Fig 5A, right) regimes to more cleanly isolate the plastic effects of each regime without directly perturbing network dynamics. Fig 5B shows the change in recall performance (Post Sleep – Pre Sleep) for the natural SWS control (gray) condition, the plasticity block during slow wave (blue) condition, and the plasticity block during delta wave (magenta) condition. Our model reproduced the prior empirical findings (see Fig 1D). Blocking plasticity during slow waves (Fig 5B; blue) resulted in a large drop in recall performance. Conversely, blocking plasticity during delta waves (Fig 5B; magenta) resulted in a substantial performance improvement beyond that of the control condition. This once again suggested that slow waves played the expected role by contributing active consolidation of the sequence memory, while delta waves were contributing to its erasure.

**Fig 5.**
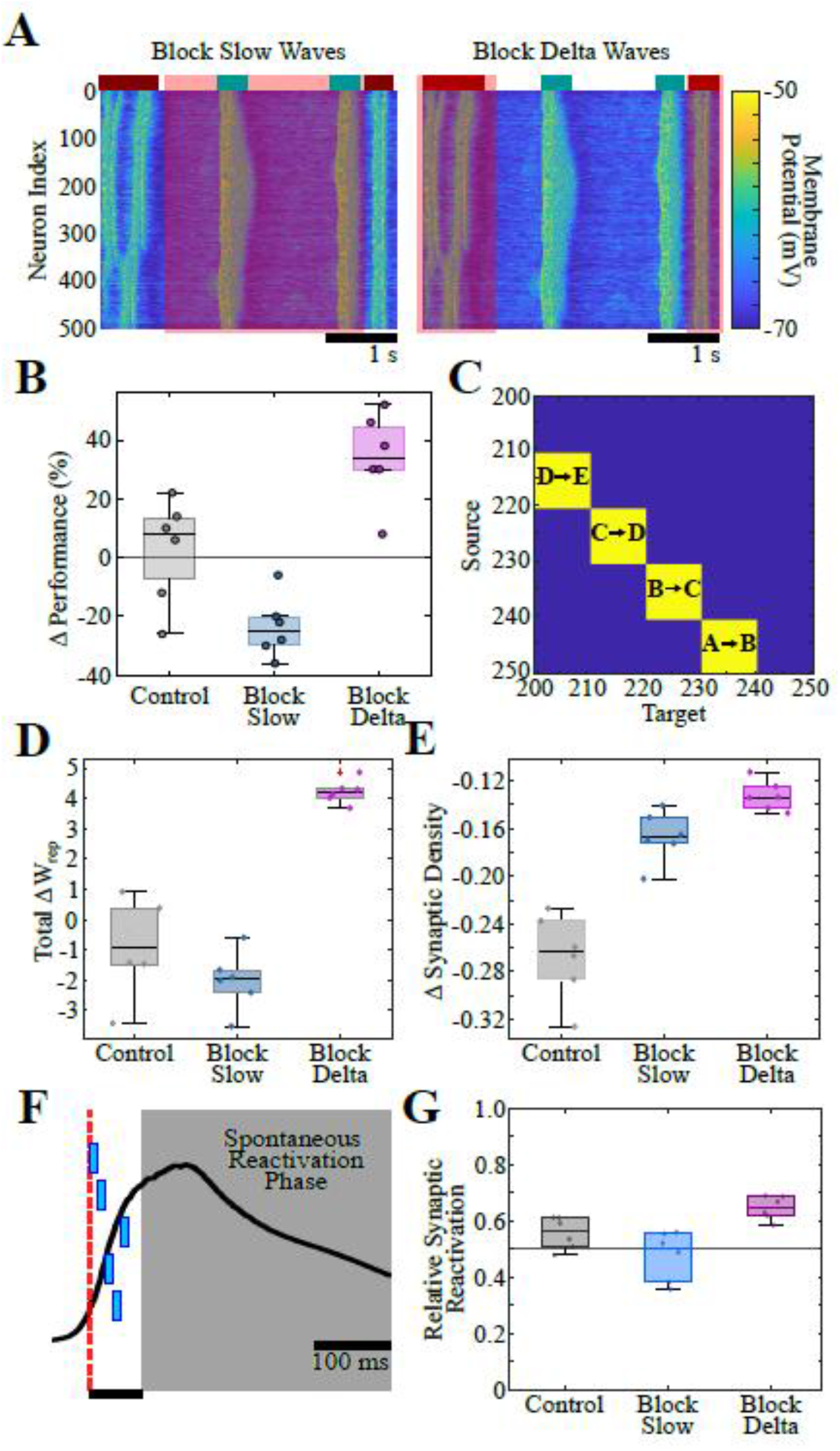
Cortical memories cannot spontaneously reactivate during delta waves. **A**. Illustration of when plasticity is blocked during slow (left) or delta (right) waves during SWS. **B**. Change in recall performance across SWS in 3 conditions: control (gray), block plasticity during slow waves (blue), and block plasticity during delta waves (magenta). **C**. Colormap showing filter used to select synapses which support the trained sequence memory, with the relevant letter group transitions inscribed. **D**. Total change in synaptic weight supporting the memory representation across SWS in 3 conditions: control (gray), block plasticity during slow waves (blue), and block plasticity during delta waves (magenta). **E**. Total change in the density of the synaptic representation across SWS in 3 conditions: control (gray), block plasticity during slow waves (blue), and block plasticity during delta waves (magenta). **F**. Schematic illustration of how when spontaneous reaction is measured during each Up state after the application of the simulated SWR input has ceased. **G**. Relative synaptic reactivation of synapses supporting the sequence memory representation across SWS in 3 conditions: control (gray), block plasticity during slow waves (blue), and block plasticity during delta waves (magenta).

Next, we analyzed how the synaptic weights supporting the memory representation (Fig 5C) were affected by each type of Up state during SWS. The total change in weight supporting the memory representation across SWS (total ΔW_rep_; Fig5D) tended to slightly decrease in the control condition (gray), it decreased dramatically when plasticity was blocked during slow waves (blue), and it increased dramatically when plasticity was blocked during delta waves (magenta). Thus, slow waves tended to increase total synaptic weight while delta waves decreased it.

To quantify the density of synaptic representation, we adopted a measure originally developed to compute the sparseness of neural population codes (Treves and Rolls, 1994) (see **Methods** for more details). Low values indicate that information is carried by a few synapses (low-density synaptic representation), whereas values near 1 indicate that information is distributed broadly across synapses (high-density representation). While SWS resulted in a lower density representation across all 3 conditions (Fig 5E), we found that it decreased the most with a mixture of slow and delta waves (gray) and the least when delta waves were blocked (magenta). Thus, delta waves were found to have a greater sparsening affect on the memory representation than slow waves.

Finally, we looked at spontaneous reactivation of synapses supporting the memory representation during the phase of each Up state after the index was applied (Fig 5F). The relative synaptic reactivation is shown in Fig 5G, which is the proportion of pre->post events divided by the total number of pre->post and post->pre spiking events, averaged across Up states. For the conditions in which plasticity was blocked, only Up states with active plasticity were considered. We found a small tendency towards spontaneous reactivation supporting the sequence memory in the control condition (gray), no tendency to reactivate during delta waves when plasticity was blocked during slow waves (blue), and a significant tendency to reactivate during slow waves when plasticity was blocked during delta waves (magenta). Thus, slow waves engage in spontaneous memory reactivation to protect it from interference and allow consolidation, but delta waves cannot.

### Extremes of Slow and Delta Regimes Reveal Asymmetric Sensitivity to Perturbation

Lastly, we utilized the flexibility of the model to explore how the probability of indexing perturbations and the ratio of slow to delta waves affected consolidation. The heatmap in Fig 6A shows the mean change in performance when the probability of indexing perturbation was reduced from 100% to 70% independently for slow and delta waves. In general, we find that as the probability of perturbation is reduced, recall performance increases from consolidation.

**Fig 6.**
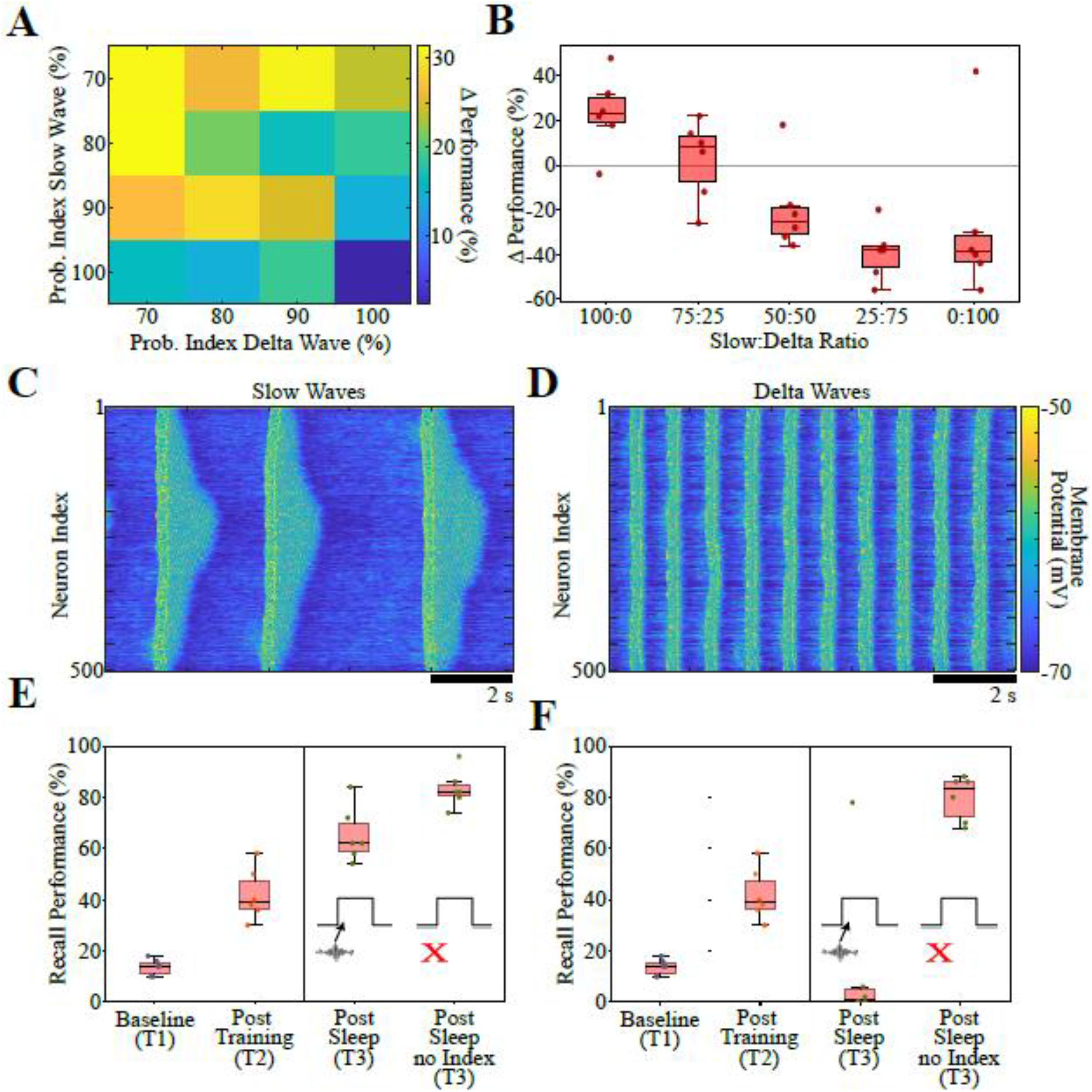
Slow and delta waves have asymmetric sensitivity to perturbation. **A**. Heatmap showing the mean change in performance across SWS when the probability of SWR input varies across slow and delta wave regimes. **B**. Boxplots showing the change in performance across SWS at the relative slow:delta wave regime ratio was systemically changed. **C**. Heatmap of PY activity during 10 s of SWS spent entirely in the slow wave regime. **D**. Same as (C) but spent entirely in the delta wave regime. **E**. Boxplots showing mean recall performance at each test period and for cases of SWR input during slow waves or not. **F**. Same as (E) but for delta waves.

Next, we varied the ratio of time spent in the slow wave compared to delta wave regime. Note that the baseline case considered thus far used a ratio of 75:25 slow:delta waves. Fig 6B shows that the extreme of 100% slow waves leads to the greatest performance gains from SWS, while increasing the ratio of delta waves results in reduced performance gains until the memory was completely forgotten by 75% delta waves. Thus, the ratio of slow:delta waves can potentially be used to control the extent of consolidation and forgetting.

Finally, we used the model to simulate SWS with only slow waves (Fig 6C) or only delta waves (6D), and either applied an indexing perturbation on every Up state or none of them. In the model with only slow waves, we found that performance increased substantially after sleep but increased greater when there were no indexing perturbations to interfere with consolidation (Fig 6E). However, in the model with only delta waves, we found that the memory is completely forgotten when indexing perturbations are applied but consolidated to the same extent as during slow waves when indexing perturbations are not applied (Fig 6F). Therefore, both slow and delta waves can potentially reactivate a memory. However, only slow waves can do this while contending with interfering input from other memories.

## DISCUSSION

In this work we implemented a model of slow and delta waves during sleep and reproduced a recent empirical phenomenon suggesting that slow waves actively consolidate memory traves while delta waves allow them to be forgotten. Moreover, the model predicted that a mixture of delta waves co-occurring with slow waves leads to sparser synaptic representation of the memory than either type of Up state on its own. Finally, we found that while both types of Up state could consolidate a sequence memory when there was no interference, only slow waves could do so when contending with interfering perturbations due to the ability to engage spontaneous reactivation later in the Up state after the perturbation. Therefore, our model predicts that the relative amount of consolidation and forgetting may be flexibly controlled by adjusting both the ratio of slow waves to delta waves, and the density of SWR input from the hippocampus during SWS.

Finally, although the two manipulations we made to the model to switch between slow and delta wave regimes were implemented as independent parameter changes, they may reflect a common physiological state. The accelerated recovery of mEPSP drive may reflect a glial-regulated synaptic state. Astrocytic Ca^2+^ signaling can modulate neuronal excitability and synaptic transmission (Goenaga et al. 2023), and astrocytic Ca^2+^ elevations have been shown to increase the frequency of miniature or spontaneous postsynaptic currents (Araque et al., 1998; Fiacco and McCarthy, 2004). Because spontaneous neurotransmitter release is itself Ca^2+^ sensitive, and astrocytic Ca^2+^ signaling varies across sleep-wake states (Bojarskaite et al., 2020), we hypothesize that delta waves may arise from a coordinated astrocyte-mediated shift in both postsynaptic Ca^2+^-dependent adaptation and increased spontaneous glutamatergic drive. While our results are largely agnostic to the particular mechanism which coordinates this shift in average Up state duration, and we can see potential mechanisms involving neuromodulators such as norepinephrine as well, we propose that astrocytic Ca^2+^ signaling may be potential unifying mechanism worthy of further investigation.

## METHODS

### Network architecture

Throughout this study, we make use of a modified version of a thalamocortical network which has been previously described in detail (Wei et al., 2016; Gonzalez et al., 2020). In brief, the network consisted of a cortical module containing 500 excitatory pyramidal neurons (PYs) and 100 inhibitory interneurons (INs), and a thalamic module containing 100 excitatory thalamocortical neurons (TCs) and 100 inhibitory reticular interneurons (REs). Connectivity in the network was determined by cell type and a local radius (see Fig. 1), and excitatory synapses were mediated by AMPA and/or NMDA currents, while inhibitory synapses were mediated by GABA_A_ and/or GABA_B_ currents.

In the cortex, PYs synapsed onto PYs and INs with a radii of R_AMPA(PY-PY)_ = 20, R_NMDA(PY-PY)_ = 5, R_AMPA(PY-IN)_ = 1, and R_NMDA(PY-IN)_ = 1. All connections were deterministic within these radii, expect for AMPA synapses between PYs, which had a 60% probability of connection. Additionally, INs synapsed onto PYs with a radius of R_GABA-A(IN-PY)_ = 5. In the thalamus, TCs synapsed onto REs with a radius of R_AMPA(TC-RE)_ = 8 and REs synapsed onto REs and TCs with radii of R_GABA-A(RE-RE)_ = 5, R_GABA-A(RE-TC)_ = 8, and R_GABA-B(RE-TC)_ = 8. Between the cortex and thalamus, TCs synapsed onto PYs and INs with radii of R_AMPA(TC-PY)_ = 15, R_AMPA(TC-IN)_ = 3, while PYs synapsed onto TCs and REs with radii of R_AMPA(PY-TC)_ = 10, and R_AMPA(PY-RE)_ = 8.

### Wake – Sleep transitions

To model the state transitions between awake and N3 sleep, we modulated the intrinsic and synaptic currents of our neuron models to account for differing concentrations of neuromodulators that partially govern these arousal state transitions. As these mechanisms have been described in detail in (Krishnan et al., 2016), here we will simply outline the approach. The model included the effects of changing acetylcholine (ACh), histamine (HA), and GABA concentrations as follows: ACh – by modulating the potassium leak current in all cell types, as well as excitatory AMPA synapses within the cortex; HA – by modulating the hyperpolarization-activated cation current in TC cells; and GABA – by modulating inhibitory GABAergic synapses within the cortex and thalamus. To transition the network from awake to sleep, we modeled the effects of reduced ACh and HA but increased GABA concentrations to reflect experimental observations (Vanini et al., 2011).

### Intrinsic currents

All cell types were modeled using the Hodgkin-Huxley formalism, and cortical PYs and INs contained dendritic and axo-somatic compartments that have been previously described (Wei et al., 2018). The dynamics of the membrane potential were modeled according to:

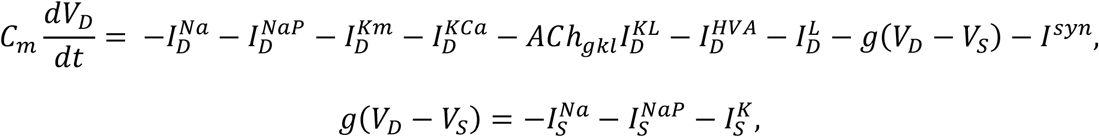

where *C*_*m*_ is the membrane capacitance, *V*_*D*_,_*S*_ are the dendritic and axo-somatic membrane voltages respectively, *I*^*Na*^ is the fast sodium (Na^+^) current, *I*^*NaP*^ is the persistent Na^+^ current, *I*^*Km*^ is the slow voltage-dependent non-inactivating potassium (K^+^) current, *I*^*KCa*^ is the slow calcium (Ca^2+^)-dependent K^+^ current, *ACh*_*gkl*_ represents the change in K^+^ leak current *I*^*KL*^ which is dependent on the level of ACh during the different arousal states, *I*^*HVA*^ is the high-threshold Ca^2+^ current, *I*^*L*^ is the chloride (Cl^−^) leak current, *g* is the conductance between the dendritic and axo-somatic compartments, and *I*^*syn*^ is the total synaptic current input to the neuron. IN neurons contained all intrinsic currents present in PY with the exception of the *I*^*NaP*^. All intrinsic ionic currents (*I*^*j*^) were modeled in a similar form:

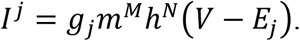

where *g*_*j*_ is the maximum conductance, *m* (activation) and *h* (inactivation) are the gating variables, *V* is the voltage of the compartment, and *E*_*j*_ is the reversal potential of the ionic current. The gating variable dynamics are described as follows:

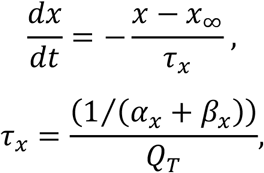

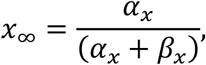

where *x* = *m* or *h, τ* is the time constant, *Q*_*T*_ is the temperature related term, *Q*_*T*_ = *Q*^((*T*−23)/10)^ = 2.9529, with *Q* = 2.3 and *T* = 36.

In the thalamus, TCs and REs contained a single compartment with membrane potential dynamics given by:

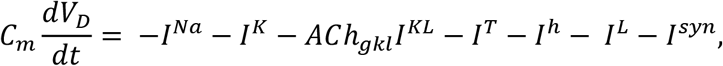

where *I*^*Na*^ is the fast Na^+^ current, *I*^*K*^ is the fast K^+^ current, *I*^*KL*^ is the K^+^ leak current, *I*^*T*^ is the low-threshold Ca^2+^ current, *I*^*h*^ is the hyperpolarization-activated mixed cation current, *I*^*L*^ is the Cl^−^ leak current, and *I*^*syn*^ is the total synaptic current input to the neurons. The *I*^*h*^ current was only expressed in TCs. The influence of histamine (HA) on *I*^*h*^ was implemented as a shift in the activation curve by *HA*_*gh*_ as described by:

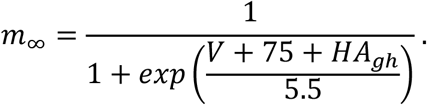

### Synaptic currents

The equations for our synaptic current models have been described in detail in our previous studies (Krishnan et al., 2016; Wei et al., 2018). To model the effects of ACh and GABA, we modified the standard equations as follows:

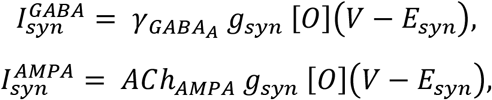

where *g*_*syn*_ is the maximal conductance at the synapse, [*O*] is the fraction of open channels, and *E*_*syn*_ is the channel reversal potential (E_GABA-A_ = −70 mV, E_AMPA_ = 0 mV, and E_NMDA_ = 0 mv). The parameter 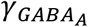 modulated the GABA synaptic currents for IN-PY, RE-RE, and RE-TC connections. For INs 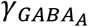 was 0.22 and 0.44 for awake and N3 sleep, respectively, while for REs 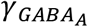 was 0.6 and 1.2. *ACh*_*AMPA*_ defined the influence of ACh levels on AMPA synaptic currents for PY-PY, TC-PY, and TC-IN. For PYs *ACh*_*AMPA*_ was 0.133 and 0.4332 for awake and N3 sleep, respectively, while for TCs *ACh*_*AMPA*_ was 0.6 and 1.2.

In addition to spike-triggered post-synaptic potentials (PSPs), spontaneous miniature PSPs (mPSPs) were implemented for both excitatory and inhibitory synapses within the cortex. The dynamics are similar to the typical PSPs described above, but the arrival times were governed by an inhomogeneous Poisson process where the next release time *t*_*release*_ is given by:

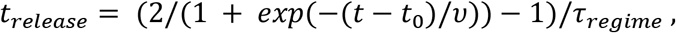

where *t*_0_ is the time of the last presynaptic spike, and *υ* was the mPSP frequency (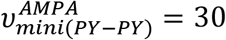, 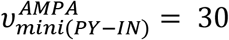, and 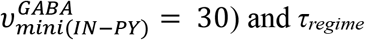) was a time constant which controlled the timescale of mPSP build-up following the last spike. The maximum conductances for mPSPs were 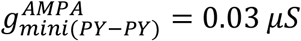, 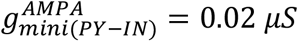, and 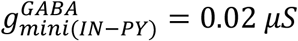.

Finally, short-term synaptic depression was also implemented in AMPA synapses within the cortex. To model this phenomenon, the maximum synaptic conductance was multiplied by a depression variable (*D* ≤ 1), which represents the amount of available “synaptic resources” as described in (Bazhenov et al., 2002). This short-term depression was modeled as follows:

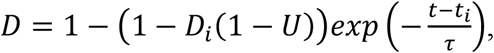

where *D*_*i*_ is the value of *D* immediately before the *i*_*th*_ event, (*t* − *t*_*i*_) is the time after the *i* _*th*_ event, *U* = 0.073 is the fraction of synaptic resources used per action potential, and *τ* = 700*ms* is time constant of recovery of synaptic resources.

### Spike-timing-dependent plasticity

The potentiation and depression of AMPA synapses between PYs were governed by the following spike-timing-dependent plasticity (STDP) rule:

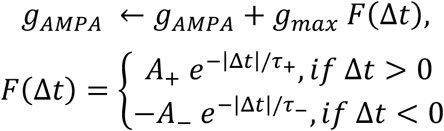

where *g*_*max*_ was the maximal conductance of *g*_*AMPA*_, *F* was the STDP kernel, and □*t* was the relative timing of the pre- and post-synaptic spikes. The maximum potentiation/depression were set to A_+/−_ = 0.002, while the time constants were set to t_+/−_ = 20 ms. A_−_ was reduced to 0.001 during training to reflect the effects of changes in acetylcholine concentration during focused attention on synaptic depression during task learning observed experimentally (Blokland, 1995; Shinoe et al., 2005; Sugisaki et al., 2016).

### Sequence training and testing

Training and testing of memory sequences was performed similarly to our previous study (Wei et al., 2018). In brief, each sequence was comprised of the same 5 groups of 10 PYs (i.e PYs 200 - 249), with Sequence 1 (S1) ordered E(240-249), D(230-239), C(220-229), B(210-119), A(200-209), and Sequence 2 (S2) ordered A(200-209), B(210-219), C(220-229), D(230-239), E(240-249). Each training bout consisted of sequentially activating each group via a 10 ms direct current pulse with a 5 ms delay between group activations. Training bouts occurred every 1 s during the training period. This training structure was chosen to ensure strong interference between S1 and S2 according to our STDP rule. Test bouts occurred every 1 ms during testing periods, in which only the first group in each sequence was activated (E for S1; A for S2), and recall performance was measured based on the extent of pattern completion for the remainder of the sequence within a 350 ms window.

### Data Analysis

All analyses were performed with standard MatLab and Python functions. For each experiment a total of 6 simulations with different random seeds were used for statistical analysis.

### Sequence performance measure

A detailed description of the performance measure used during testing can be found in (Wei et al., 2018) and the code is available in (https://github.com/o2gonzalez/sequencePerformanceAnalysis) (Gonzalez et al., 2020). Briefly, the performance of the network for recalling a given sequence following activation of the first group of that sequence was measured by the percent of successful sequence recalls. We first detected all spikes within the predefined 350 ms time window for all 5 groups of neurons in a sequence. The firing rate of each group was then smoothed by convolving the average instantaneous firing rate of the group’s 10 neurons with a Gaussian kernel with window size of 50 ms. We then sorted the peaks of the smoothed firing rates during the 350 ms window to determine the ordering of group activations. Next, we applied a string match (SM) method to determine the similarity between the detected sequences and an ideal sequence (ie. A-B-C-D-E for S1). SM was calculated using the following equation:

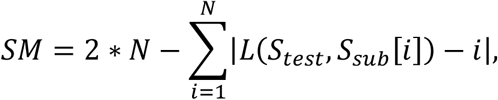

where *N* is the sequence length of *S*_*test*_, *S*_*test*_ is the test sequence generated by the network during testing, *S*_*sub*_ is a subset of the ideal sequence that only contains the same elements of *S*_*test*_, and *L*(*S*_*test*_, *S*_*sub*_[*i*]) is the location of the element *S*_*sub*_[*i*] in sequence *S*_*test*_. *SM* was then normalized by double the length of the ideal sequence. Finally, the performance was calculated as the percent of recalled sequences with *SM* ≥ *Th* = 0.8, where *Th* is a threshold indicating that the recalled sequence must be at least 80% similar to the ideal sequence to be counted as a successful recall as previously done in (Wei et al., 2018).

### Density of synaptic weight states

In order to compute the density of synaptic weight matrices, we adapted a measure previously conceived to compute the sparsity of neural population codes, operating over real-valued firing rates (Treves and Rolls, 1994),

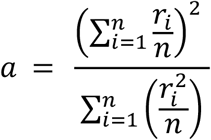

where *a* is the density of the population code, *r*_*i*_ is the firing rate of the *i*th neuron, and *n* is the total number of neurons. The density here is maximal at 1 when all neurons have the same firing rate, 1/*n* when only one neuron responds, and zero when the population is silent.

To adapt this to compute the density of synaptic weight states, we first unrolled the synaptic weight matrix into a vector and then removed all non-existent synapses (i.e. zero-valued) from the vector so that we only computed over the number of existing synapses rather than potential synapses. We then computed

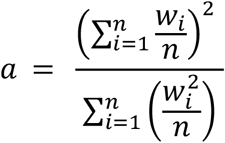

where *a* is the density of the weight state, *w*_*i*_ is the firing rate of the *i*th synapse, and *n* is the total number of synapses. Thus, the density here is maximal at 1 when all synapses have the same strength, and near 1/*n* when only one synapse is above its minimum strength value (note that existing synapses could not drop to zero strength).

### Online DUt and UDt detection

Online detection of the DUt and UDt during the simulations was conducted by tracking the simulated global LFP in the same manner as reported previously ^23^. Briefly, the LFP was approximated by calculating mean membrane potentials of all the cortical excitatory neurons, and it had a bimodal distribution during N3 sleep, where one peak corresponded to the Up state and another peak to the Down state. The trough of the distribution was selected as a threshold to separate Up and Down state. The onset of Up or Down state was then defined as the moment when LFP value crossed the threshold.

### Synaptic reactivation analysis

First, each Up state was identified by detecting the DUt and UDt times during SWS for a given simulation. Each Up state was then divided into an Indexed Phase which began at the application of the index (i.e. the DUt) and lasted for 80 ms, and a Spontaneous Reactivation Phase which began at the end of the Indexed Phase and lasted until the end of the Up state (i.e. the UDt).

Next, focusing on a single Up state, the PY spike times during the Spontaneous Reactivation Phase were inserted into an offline STDP implementation which was identical to the one used online in simulation. Using this implementation, for a given pair of synaptically connected PYs, we computed all pre->post and post->pre events which occurred during this phase of the Up state, summed these event types, and computed the relative proportion of pre->post events as follows:

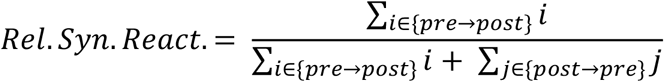

